# Preliminary Study of the Potential Mechanism of CTSD in the Aging Process of *Sepiella japonica*: Fundamental Function Analysis

**DOI:** 10.1101/392688

**Authors:** Baoying Guo, Yu Chen, Pengzhi Qi, Yingying Ye, Zhenming Lv, Xiumei Zhang, Kaida Xu

**Author notes:** Corresponding author: Baoying Guo, E-mail address (B.Y. Guo).

## Abstract

Cathepsin D, a kind of endopeptidase, can degrade peptides and proteins in lysosomes, which are involved in cell apoptosis. Previous transcriptome analysis of optic glands of *Sepiella japonica* across four growth stages, expression of a cathepsin D-like segment was found to be significantly different. Based on the complete cDNA sequence of *S. japonica*, the CTSD gene (also called sjCTSD, GenBank accession no. KY745896.1) was cloned using RACE amplification; this gene is 1389 bp in length and encodes proteins composed of 393 amino acids. Spatio-temporal expression profiles of the sjCTSD gene were determined using qPCR assays, which showed that the expression levels of sjCTSD constantly increased across four growth stages in 9 of 11 tissues that were investigated. In the optic glands, as well as pancreas and liver cells, sjCTSD expression levels sharply increased during the post-spawning phase. To investigate the potential role of the sjCTSD gene in aging progress, we constructed the prokaryotic expression vector of pET28a/sjCTSD. After induced by IPTG, the recombinant protein sjCTSD was obtained in the form of inclusion bodies, with a molecular size of approximately 40.3 kDa, the inclusion body of sjCTSD can be converted to soluble protein through the denaturation and renaturation. The results of functional experiments showed that sjCTSD could degrade bovine hemoglobin under acidic conditions, and inhibit the growth of *Escherichia coli* and *Vibrio alginolyticus*, which was speculated that the increased expression of sjCTSD may help inhibit the invasion of pathogenic bacteria with the immune function of cuttlefish declines during the aging process. To a certain extent, these results indicated the potential functional role of the sjCTSD gene in the aging process of *S. japonica*. This study provides insights to further understand the roles of lysosomal proteins on anti-aging effects in *S. japonica* and other cephalopoda species.

## 1 Introduction

*Sepiella japonica* (Sepiidae, Sepiella) is endemic to the coastal waters of Eastern Russia, Japan, North Korea and China. It is the only member of the genus *Sepiella* found in the China sea and an economically valuable fishery resource and considered one of the four most famous sea products in China mainly due to highly edible, medicinal and economic value (Shen et al., 2007). However, the wild population has dramatically declined since the 1980s and has become nearly extinct, presumably due to over exploitation, environmental disruption and in particular, the deterioration of the spawning grounds (Wu et al., 2006). The cuttlefish is annual, the brood amount is only 1800, much less than that of fishes or shrimps, which are about 200,000 and even 1,000,000, even more regrettably, it has been reported that a large number of eggs (approximately 60% of brood amount) remained in the abdomen of naturally dead female *S. japonica*, indicating immense loss of reproductive resource. However, why this phenomenon happen remains unknown, as very interesting and important biological topic, which requires urgent attention from cephalopod researchers to determine the possible reason and propose solutions subsequently. If the death or aging process of *S. japonica* could be postponed artificially, more offspring would be obtained for marine aquaculture or proliferation, which will greatly help to recovery this precious marine resources.

Aging is an inevitable physiological change occurring in organisms over time (Bowen et al. 2004), which ultimately leads to death naturally, as one gets old, along with gradual dysfunctions of all organs in the organisms including unicellular organisms, plants, animals and humans. The growth and aging of the body can not be separated from the regulation of cell apoptosis, according to the type of protein hydrolysis, cathepsin can be divided into cysteine protease, aspartic acid protease, metalloprotease, serine protease, glutamic acid protease and threonine protease (Repnik et al. 2012). Aspartic protease plays a key important role in the process of regulating cell apoptosis to mediate tumor cell death (Ivanova et al. 2008; Vasiljeva et al. 2008; Dong et al. 2012; Kang et al. 2017).

Cathepsin D (CTSD), a type of endoproteolytic aspartic proteinase, plays a key role in lysosomal digestive activity (Tsuji et al., 1994; Dell’Angelica et al., 2000; Feng et al., 2011). CTSD activity was found to be enhanced in wide variety of tissues during remodeling, regression, cell proliferation and apoptosis (Zuzarte-Luis et al., 2007; Mei et al., 2008; Gonçalves et al., 2016; Reddy et al., 2017). Cathepsin D has received much attentions for its potential importance in aquaculture (Kenessey et al., 1989; Cheret et al., 2007; Pan et al; 2012), but few studies have been conducted on the function of CTSD in the aging process (Kenessey et al., 1989; Cheret et al., 2007; Pan et al., 2012).

Furthermore, our lab transcriptome data (not published) of four growth periods (young, pre-spawning after sexual maturity, spawning, and postspawning before death) indicated that the fragment marked as CTSD was significantly differentially expressed genes. To further investigate the potential role of CTSD in the growth and aging of *S. japonica*, we analyzed the mRNA levels of CTSD in different tissues at four growth periods, and constructed the prokaryotic expression vector of sjCTSD, attempting to find the relation between CTSD gene and aging process of S. japonica. These data could be useful to improve our understanding about CTSD functions and molecular mechanisms related to the aging process of S. japonica.

## 2 Materials and methods

### 2.1 Animals and sample collection

Individuals of *S. japonica* were collected from the Cangnan Cuttlefish Breeding Base from May to July 2016 (Ningde City, Fujian Province, China). The cuttlefish were bred under the same standard conditions and the samples were collected from four different growth phases (young, pre-spawning after sexual maturity, spawning, and postspawning before death). Eleven tissues (heart, liver, pancreas, stomach, intestines, gill, skin, mantle, brain, optic lobe, and optic gland) were dissected from each healthy cuttlefish of four growth phases. All the tissues collected were immediately preserved in liquid nitrogen and stored at - 80 °C until RNA extraction.

### 2.2 RNA isolation and cDNA synthesis

Total RNA was extracted from the tissues using an RNA extraction kit (E.Z.N.A. Total RNA KitII, OMEGA bio-tek, TaKaRa, Dalian, China), following the manufacturer’s instructions. The RNA samples were then dissolved in RNase-free water and stored at -80 °C until further analysis. The Transcription First Strand cDNA Synthesis Kit (Roche, Shanghai, China) was used to reverse transcribe RNA to cDNA, according to the manufacturer’s instructions. The reaction parameters of cDNA synthesis were as follows: the first step at 65 °C for 10 min, the second was performed at 25 °C for 10 min and 50 °C for 60 min. The cDNA was stored at -20 °C for further experiment.

### 2.3 RACE

The primer was designed according to the sequence, which was annotated as cathepsin D in the transcriptomic data of *S. japonica* (Table. 1 for partial CTSD). After sequencing validation by the LiuHe HuaDa Biotechnology Co. Ltd. (Beijing, China), the fragment was verified to be part of open reading frame of *S. japonica* cathepsin D (sjCTSD).

**Table 1.**
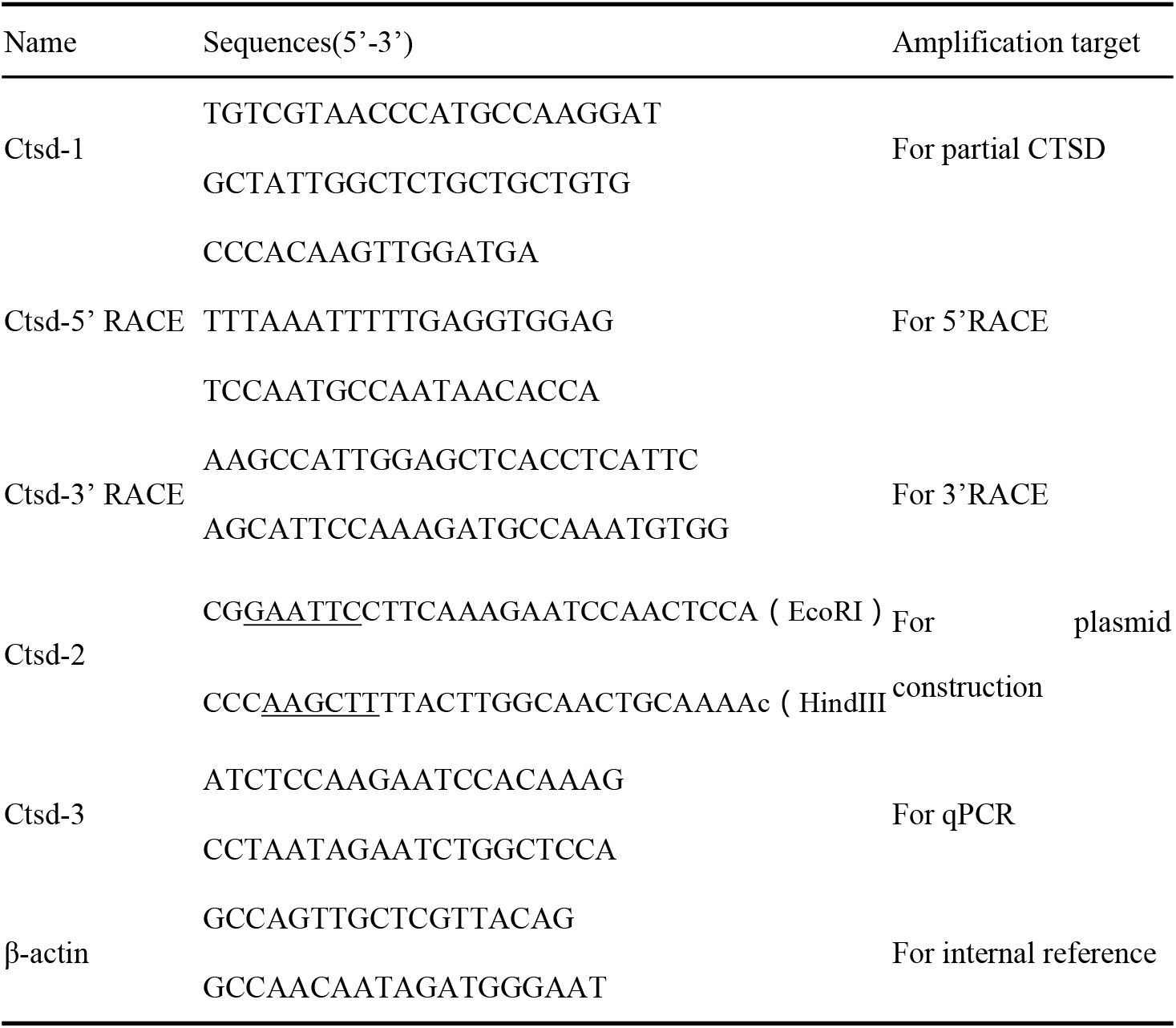
Oligonucleotide primers used for studies of CTSD gene cloning, expression and qRT-PCR.

The full-length sjCTSD cDNA sequence was obtained using rapid amplification of cDNA ends (RACE) with a 5′/3′ rapid-amplification of cDNA ends (RACE) kit (Takara Bio Inc., Dalian, China), according to the manufacturer’s protocol. Three gene-specific primers (B082-1, B082-2, B082-3) for 5′RACE and two specific primers (C063-1 and C063-2) for 3′RACE were synthesized by Shanghai Sangon Biological Engineering Technology & Services Co., Ltd. (Shanghai, China). Briefly, for the 5′-RACE reactions, we used GSP-1 primer and Superscript II RT enzyme for synthesizing cDNA. The first round of PCR was performed using GSP-2 and AAP primers, the second round of PCR was performed using GSP-3 and AUAP primers (Table 1). C063-1 and C063-2 were used for the 3’-RACE with the UPM primer. The 5′/3′ RACE products were purified, cloned, and sequenced by Shanghai Sangon Biological Engineering Technology & Services Co., Ltd.

### 2.4 Bioinformatic analysis

The CTSD mRNA sequence was analyzed using NCBI ORF Finder computational tool (http://www.ncbi.nlm.nih.gov/gorf/gorf.html). The analyses of the nucleotide and amino acid sequence homologies were performed using NCBI Blast (http://blast.ncbi.nlm.nih.gov/). The ProtScale (http://web.expasy.org/protscale/) web-based programs were used to predict the physiochemical properties of the CTSD protein. The conserved domains were predicted using the SMART online tool (http://smart.emblheidelberg.de/). Potential leader peptides were predicted using SignalP 4.0 (http://www.cbs.dtu.dk/services/SignalP/). The N-glycosylation sites, phosphorylation sites and the secondary structure of the CTSD protein were predicted by NetNGlyc 1.0 (http://www.cbs.dtu.dk/services/NetNGlyc/), Netphos 2.0 Server and HNN (http://npsa-pbil.ibcp.fr), respectively. A phylogenetic tree was constructed using the MEGA program based on the neighbor-joining method, with 1000 bootstrap replicates.

### 2.5 Expression analysis of sjCTSD by qRT-PCR

#### 2.5.1 Gene expression analysis

The expression profiling of sjCTSD mRNA in different tissues and across different growth stages were assessed using qRT-PCR. The qRT-PCR was performed in a 96-well plate (Bio-Rad Laboratories, USA) using SYBR^®^ Premix Ex Taq™ (Takara Bio Inc., Dalian, China). Briefly, the 20 μL reaction consisted of 0.4 μL of cDNA, 10 μL of SYBR^®^ Premix ExTaq™, each 0.8 μL of forward primer (10 μM), and nuclease-free water. This is the cycling paramenter of qRT-PCR, as follow: Initial denaturation at 95 °C for 10 s, repeat amplification by 40 cycles at 95 °C for 5 s, 58 °C for 30 s, and 72 °C for 30 s. The relative quantitation of gene expression was performed in three replicates for each sample. *β*-actin was considered as an internal control. The relative levels were calculated using the 2^−ΔΔCt^ method.

#### 2.5.2 Statistical analysis

The data were presented as mean ± standard deviation (S.D.) for at least three independent experiments. Statistical significance was analyzed by one-way analysis of variance or Student’s t-test using SPSS, version 16.0 (SPSS, Inc., Chicago, IL, USA). \**p*< 0.05 were considered as a statistically significant difference.

### 2.6 Plasmid construction and transfection

The primer with EcoR I and Hind III restriction sites of the sjCTSD without leader peptide was designed (Table. 1). The amplified products were gel-purified and cloned into pMD18-T vector (Takara, Dalian, China). After being transformed into *E. coli* Top5, the positive clones were identified by PCR and sequenced. Then the fragment of sjCTSD and the expression plasmid vector pET-28a(+) were digested with EcoR I and Hind III, respectively, then were ligated with T4 DNA Ligase (Takara, China). The resulting recombinant plasmid pET-28a(+)-sjCTSD was identified by PCR, enzyme cleavage and sequencing to confirm the correct insert of sjCTSD gene.

### 2.7 The expression and purification of sjCTSD protein

The recombinant plasmid pET-28a(+)-sjCTSD was transformed into *E. coli* BL21, and incubated in Luria-Bertani (LB) broth with50 μg/ml kanamycin overnight at 37 °C under shaking (200 rpm). A 100 μL aliquot of the overnight culture was added into 3 ml fresh LB medium containing kanamycin, the latter was incubated at 37 °C on 200 rpm shaker until an OD600 of approximately 0.6 was obtained, then, isopropyl β-D-thiogalactoside (IPTG) was added into the culture to a final concentration of 1 mM for the induced expression of the recombinant protein at 37 °C for 6 h. The bacterial cells were harvested and washed using 1 × phosphate buffer saline (PBS) (pH 7.4) for three times. Cells were subjected to sonication at 150 W for 20 times with 6 s working and 6 s free on ice. After centrifugation at 12,000 r/min for 5 min at 4 °C, the supernatant was stored at −20 °C, the precipitate containing inclusion body was resuspended by 1 × PBS before being preserved at −20 °C. The inclusion body undergoes a process of denaturation and renaturation, restoring its correct conformation. Then, the protein with correct conformation was purified using the 6× His-Tagged Protein Purification Kit (Cwbio, China) according to the manufacturer’s specifications. The expression of the 6× His-Tagged sjCTSD was confirmed on a 12% sodium dodecyl sulfate polyacrylamide gel electrophoresis (SDS-PAGE) and was visualized by staining with coomassie brilliant blue. The protein concentration was measured using a NanoDrop 2000 spectrophotometer with an absorption at 280 nm.

### 2.8 Enzyme activity and antibacterial of sjCTSD

In this experiment, the hemoglobin hydrolysis assay was used to detect the activity of cathepsin D. Using bovine hemoglobin as a substrate, the sjCTSD protease activity was measured at 25°C at different pH value (pH = 2.5, 3.0, 3.5 and 4.5) (the optimum culture temperature for *S. japonica* is from 24°C to 26°C). Three protease inhibitors (Pepstain A, EDTA and PMSF) were used to perform inhibition experiments on sjCTSD protein. Added 40 μL citrate buffer (pH=3.0) and 20 μL sjCTSD protein to the tube and mix and incubate for 3h at 25°C. Added 2μL inhibitor to continue incubation for 30 min to fully react the inhibitor and sjCTSD, the enzyme activity was measured every 20 minutes. After the termination of the reaction, the mixture was centrifuged (4°C, 13,000 rpm) for 20 minutes, then determined the protein concentration of the supernatant.

*E. coli* and *Vibrio alginolyticus* were selected as experimental objects for detecting bacteriostatic properties of sjCTSD. After expanding the culture strain, 10 μL of the bacterial solution was added to 200 μL of activated sjCTSD protease solution to different concentrations (50, 100, 200, 300 μg/mL) and incubated at 25°C for 2 hours. The control group used the CHAPS containing DTT instead of the sjCTSD protein solution, and the rest of the treatments were consistent. Each solid medium plate contained 20 μL of the mixture, each sample was subjected to four replicates and counted after incubation at 37° C for 10 h.

## 3 Results

### 3.1 Sequence analysis of sjCTSD

The cDNA was 1392 bp long with an ORF of 1179 bp, starting from a putative initiation codon at nucleotide 59, a 58-bp 5′ untranslated region (UTR), and a 152-bp 3′ UTR (Fig. 1). A typical polyadenylation signal sequence AATAAA was located at 15 nt upstream from the poly (A) tail. The complete ORF encoded a putative protein of 392 amino acids with a calculated molecular weight of 38 kDa (Genbank accession no. KY745896.1). A Genbank database search using BLAST revealed that the sjCTSD sequence was consistent with the sjCTSD sequences of *Todarodes pacificus*, with 83% identity. However, the predicted protein shared 80%, 68%, 67%, 66%, 65%, 64% and 62% amino acid sequence similarity with the sjCTSD protein sequences of *T. pacificus, Pinctada maxima, Pinctada margaritifera, Pteria penguin, Polyrhachis vicina, Anopheles darlingi* and *Azumapecten farreri* (Fig. 2). The alignment analysis results showed that the sjCTSD gene and putative amino acid sequences of the *S. japonica* were most similar to that of *T. pacificus*.

**Figure 1.**
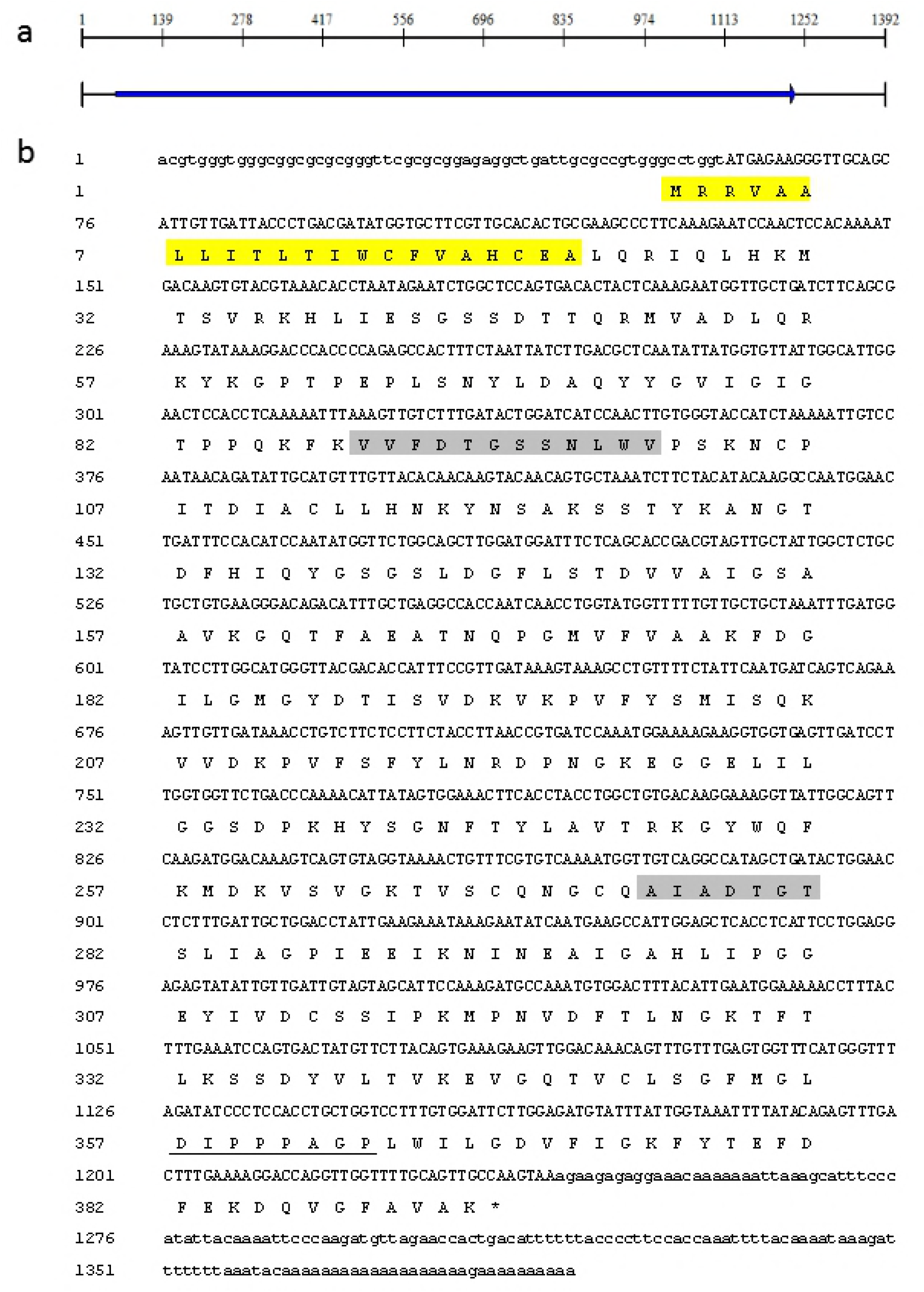
The full - length cDNA of CTSD gene and the assorted amino acid aequence of *S. japonica*. Note: * represents the terminal codons; the leader peptide sequence was labeled yellow; the gray shades were two aspartic protease signature sequences; the characteristic sequence of non-digestive cathepsin D was underlined.

**Figure 2.**
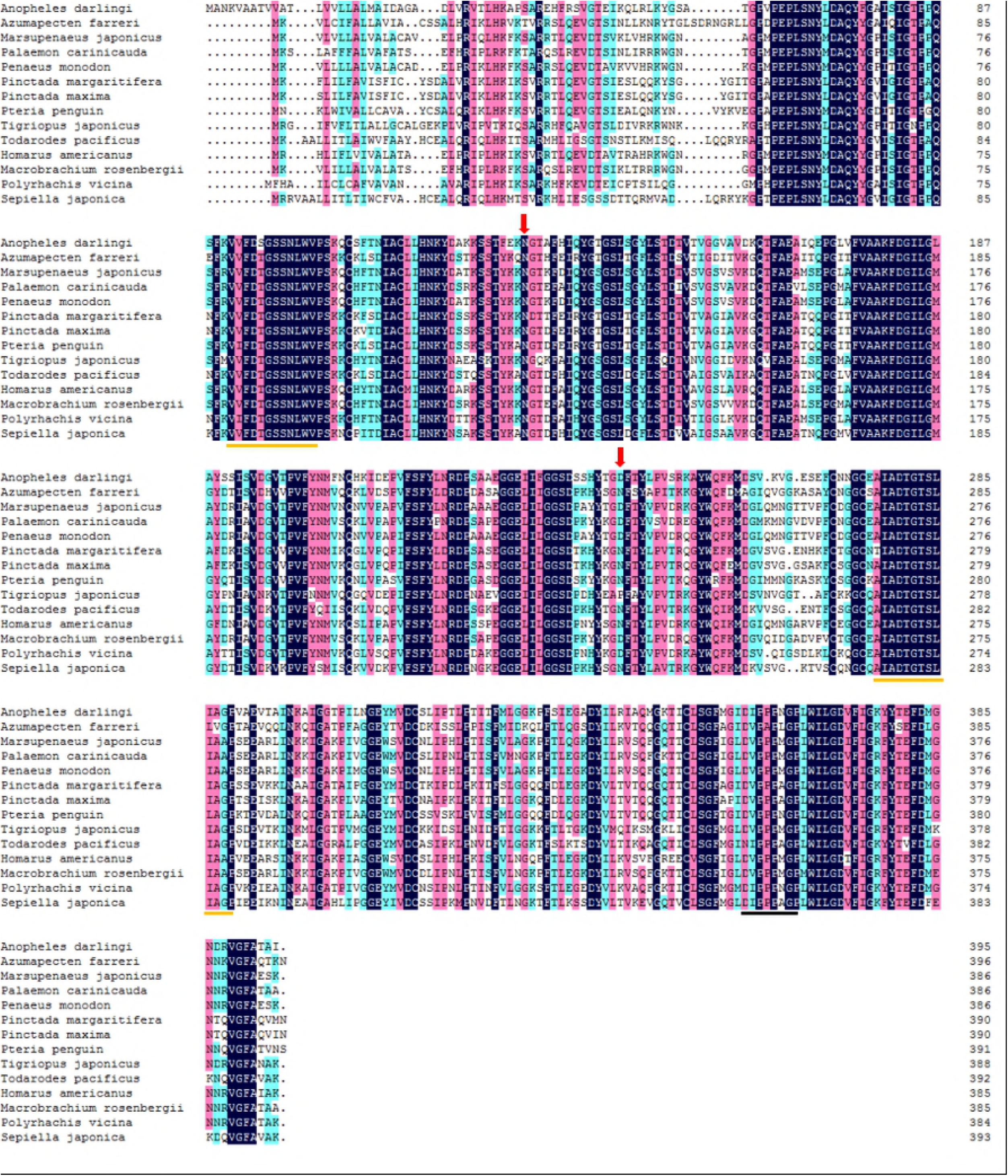
The amino acid sequence of *S. japonica* CTSD alignment with other species. Aspartic protease signature sequences were surrounded by an orange outline. Two putative N-glycosylation sites were marked by a red arrow.

### 3.2 SjCTSD protein structure prediction and phylogenetic analysis

The phylogenetic analysis showed that the *S. japonica* CTSD protein was most similar to that of *T. pacificus* (Fig. 3). The physicochemical analysis of the sjCTSD protein revealed a molecular formula of C_1934_H_3005_N_495_O_567_S_17_ and a molecular mass of 42.81 k Da. The theoretical isoelectric point of the sjCTSD protein was 4.79. The instability index of the sjCTSD protein was computed as 28.50, suggesting the protein was stable. The average hydropathicity was −0.193. The results of the online SMART tool showed that the functional domains of the sjCTSD included the leader peptide domain (Met1-Ala22), the pro-peptide domain (Leu23-Gly42) and the mature peptide domain (Ger43-Lys393). The secondary structure of the sjCTSD protein was predicted to consist of 12.21% alpha helix, 34.10% extension chain and 53.69% irregular curl. Two putative N-glycosylation sites were identified in amino acid positions 129 and 242 (Asn). In addition, *S. japonica* CTSD had two aspartyl protease active sites which were predicted from 89 to 101 aa and 275 to 287 aa, as well as two aspartic protease signature sequences: VVFDTGSSNLWV and AIADTGTSLIAG.

**Figure 3.**
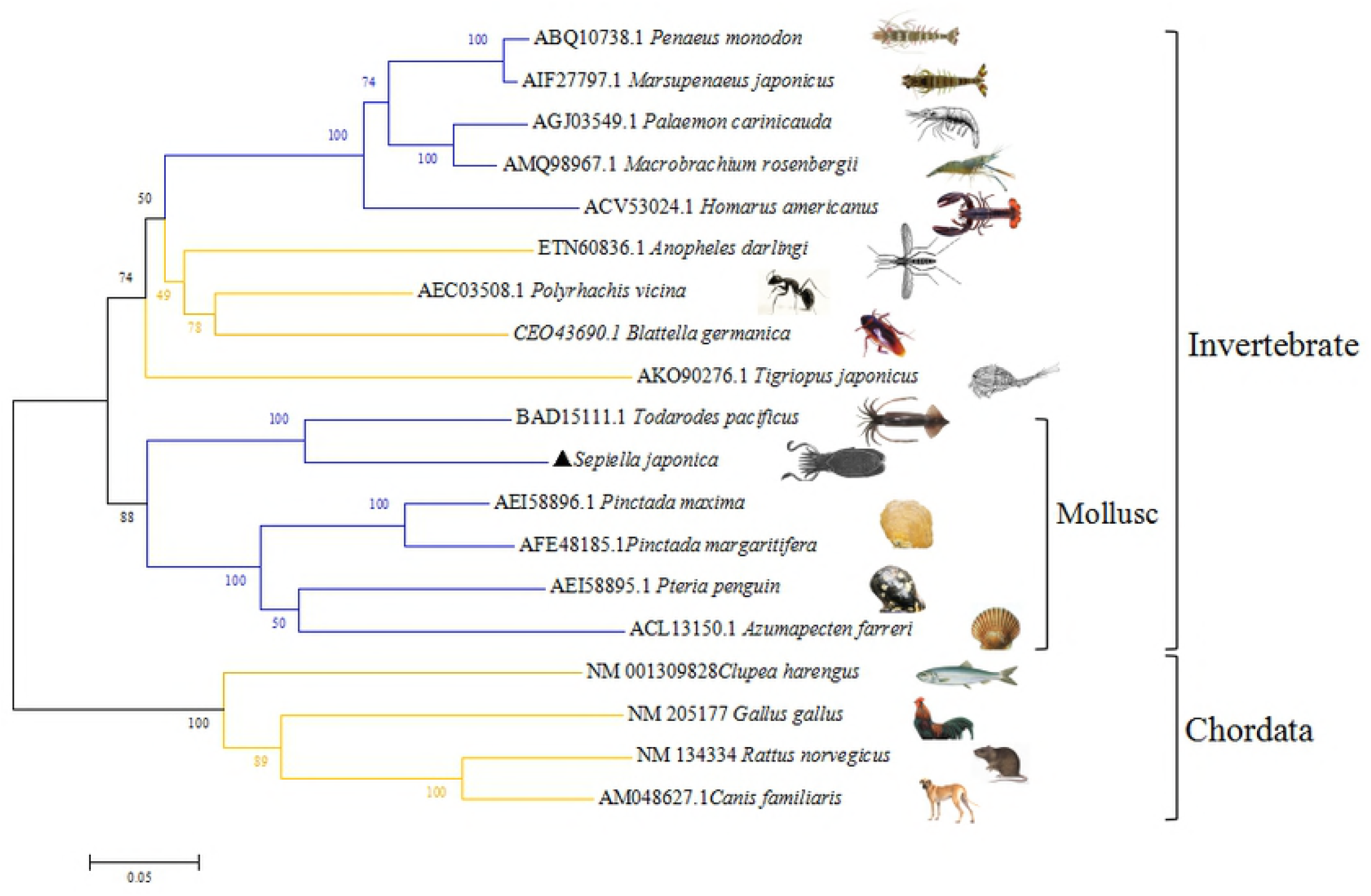
NJ Phylogenetic tree of CTSD of species. Note: The bifurcation value represents the confidence level obtained by repeated sampling of 1000 times; The ruler length represents 0.05 substitutions per site.

### 3.3 SjCTSD mRNA and protein expression levels in different tissues

RT-PCR analysis showed sjCTSD mRNA expression in all tissues (Fig. 4). At the youngest stage, the expression levels of sjCTSD gene in stomach and optic lobe tissues were higher than in other tissues, but not significantly. When the cuttlefish was at the stage of gonadal maturation, the expression level in optic lobe tissue was significantly higher than in other tissues. At the spawning stage, the expression level was significantly increased in the brain, liver, heart and optic glands (*p* < 0.05). At the stage of postspawning before death, sjCTSD mRNA expression in the optic glands, pancreas and liver were significantly higher compared with other tissues (*p* < 0.05). In contrast, the low expression level was observed in the skin, mantle and intestines (*p* < 0.05).

**Figure 4.**
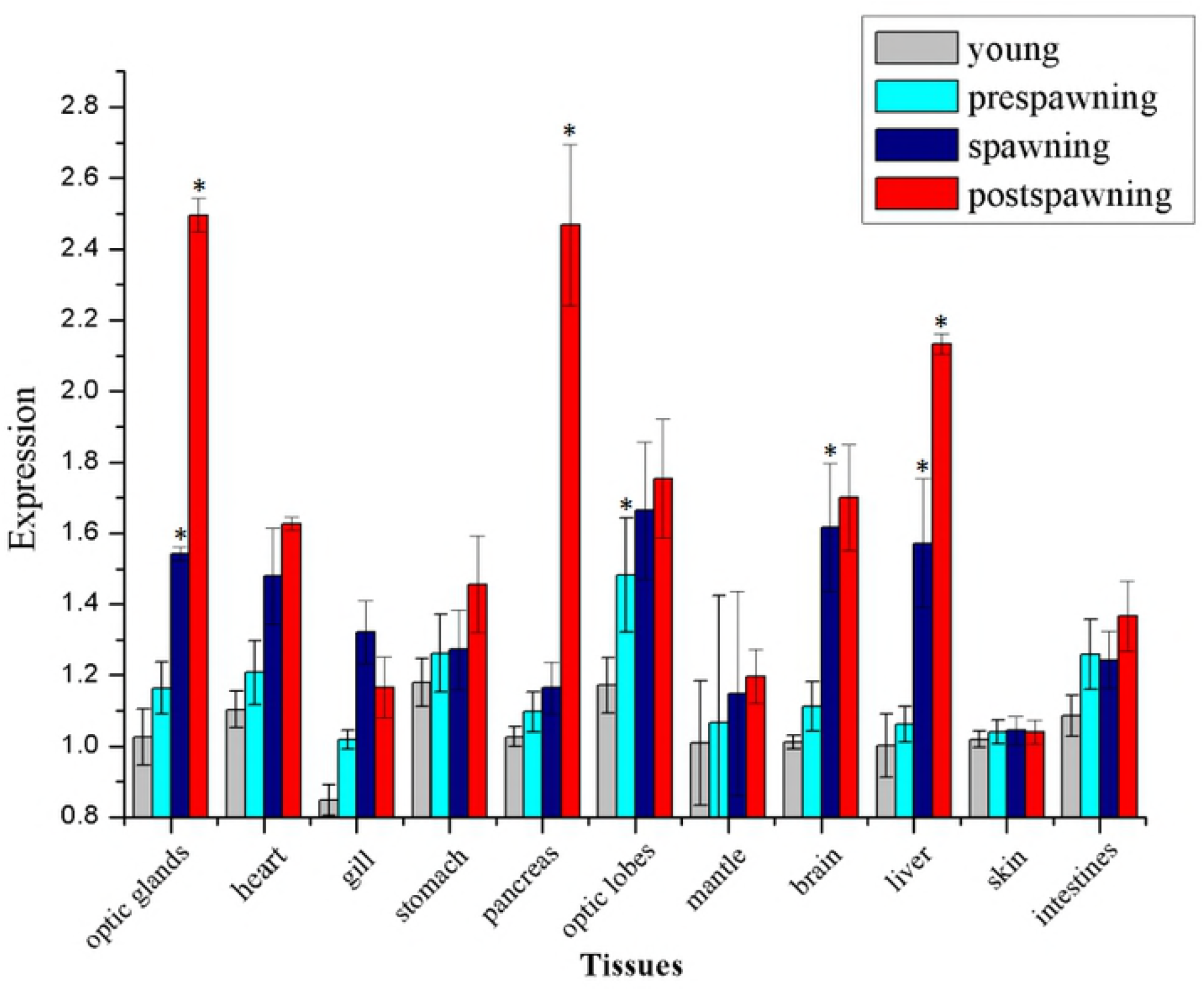
Tissue distribution of CTSD in different growth stages of *S. japonica*.

### 3.4 Expression profiling of sjCTSD mRNA within the growth stages

Expression of the sjCTSD gene in different growth stages of all tissues of *S. japonica* was monitored using qRT-PCR (Fig. 4). The transcript level of sjCTSD mRNA in the young stage was set as the control group. Between the young and prespawning stages, the expression levels were not significantly increased in any tissues (*p* > 0.05). Compared with prespawning, the level of sjCTSD expression in the spawning stage was significantly higher in the brain, optic glands and liver of the cuttlefish (*p* < 0.05). The expression level of sjCTSD in optic glands, liver and pancreatic tissues at the postspawning stage were significantly comparing to the spawning stage (*p* < 0.05). With individual growing, sjCTSD expression levels gradually increased in most tissues, except in the skin and mantle, the expression level in the skin and mantle had almost no change during the whole growth period (*p* > 0.05).

### 3.5 pET-28a(+)-sjCTSD

The primers containing EcoR I and Hind III restriction enzyme site were used to amplify ORF of sjCTSD gene. As shown in Fig. 5, sjCTSD has been ligated into the pET-28a(+) vector successfully. The proper construction of the expression plasmid pET-28a(+)-sjCTSD was identified by PCR, restriction enzyme digestion and nucleotide sequencing. The sequence contained an initiating ATG, His-tag and lac operator provided by express pET-28a(+) vector at its N-terminal.

**Figure 5.**
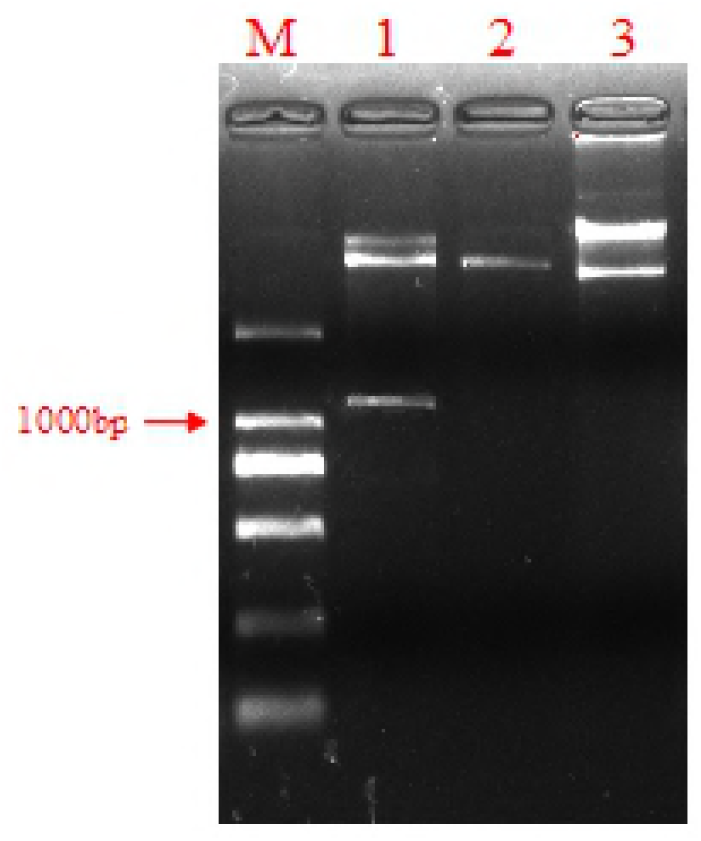
Identification of pET28a/sjCTSD recombinant plasmids by AGE (1: after double digestion of pET28a/sjCTSD recombinant plasmids; 2: double digestion of pET28a plasmids; 3: undigested pET28a/sjCTSD recombinant plasmids; M: 2000 marker)

### 3.6 Expression of sjCTSD in *E. coli*

The recombinant plasmid pET-28a(+)-sjCTSD was transfected into *E. coli* by freeze thawing methodI and induced by IPTG. The recombinant protein of sjCTSD was highly and effectively expressed in *E. coli* BL21 as a major protein product in the total cellular protein, compared to the negative control. As for the weight of sjCTSD (without leader peptide) protein, the predicted molecular weight was 40.3 kDa, and the N-terminal fusion protein with His labeling of amino acids was about 5 kDa, a total of about 45.3 kDa. Expressed recombinant proteins was in precipitation as a form of inclusion bodies by SDS-PAGE detection and the fusion protein band’s corresponding size was consistent with the expected value in SDS-PAGE (Fig. 6).

**Figure 6.**
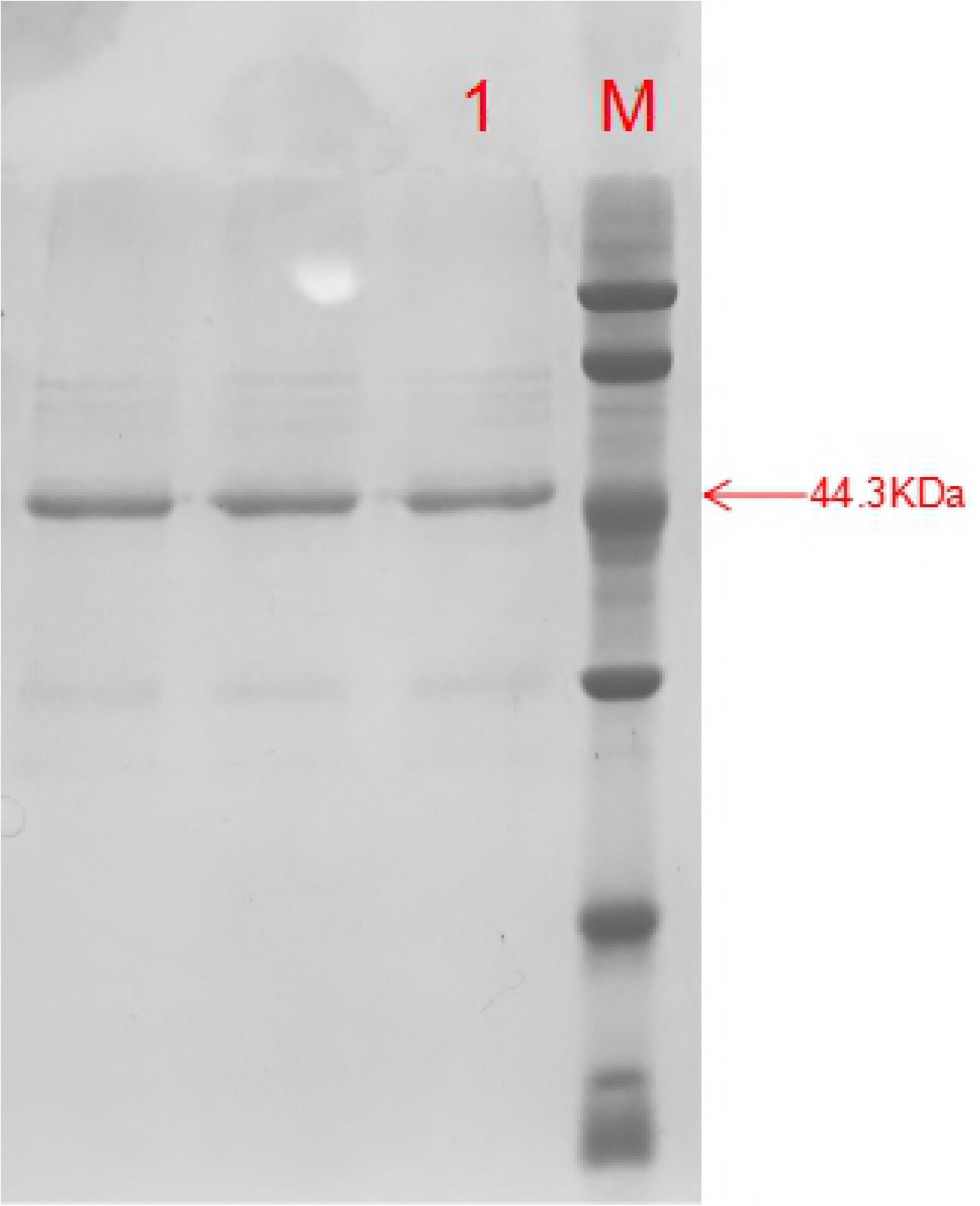
Purified sjCTSD recombinant protein. 1: sjCTSD purified protein; M: low protein Marker.

### 3.7 Enzyme activity detection of sjCTSD

Figure 7 showed that, when the pH value was between 2.5 and 3.5, the sjCTSD recombinant protein could be reactivated by self-cutting at 25°C. As the results in our study that the absorbance at A280 nm was positively correlated with the enzyme activity of CTSD. The enzyme activity of sjCTSD was the highest at pH=3.0. The enzymatic activity of recombinant sjCTSD could be inhibited by pepstain A (Fig. 8). In comparison, the inhibitory effects of EDTA and PMSF on sjCTSD enzyme activity were not notably. It was shown that the recombinant sjCTSD protein was functionally an aspartic protease and the renaturation of sjCTSD inclusion bodies was successful.

**Figure 7.**
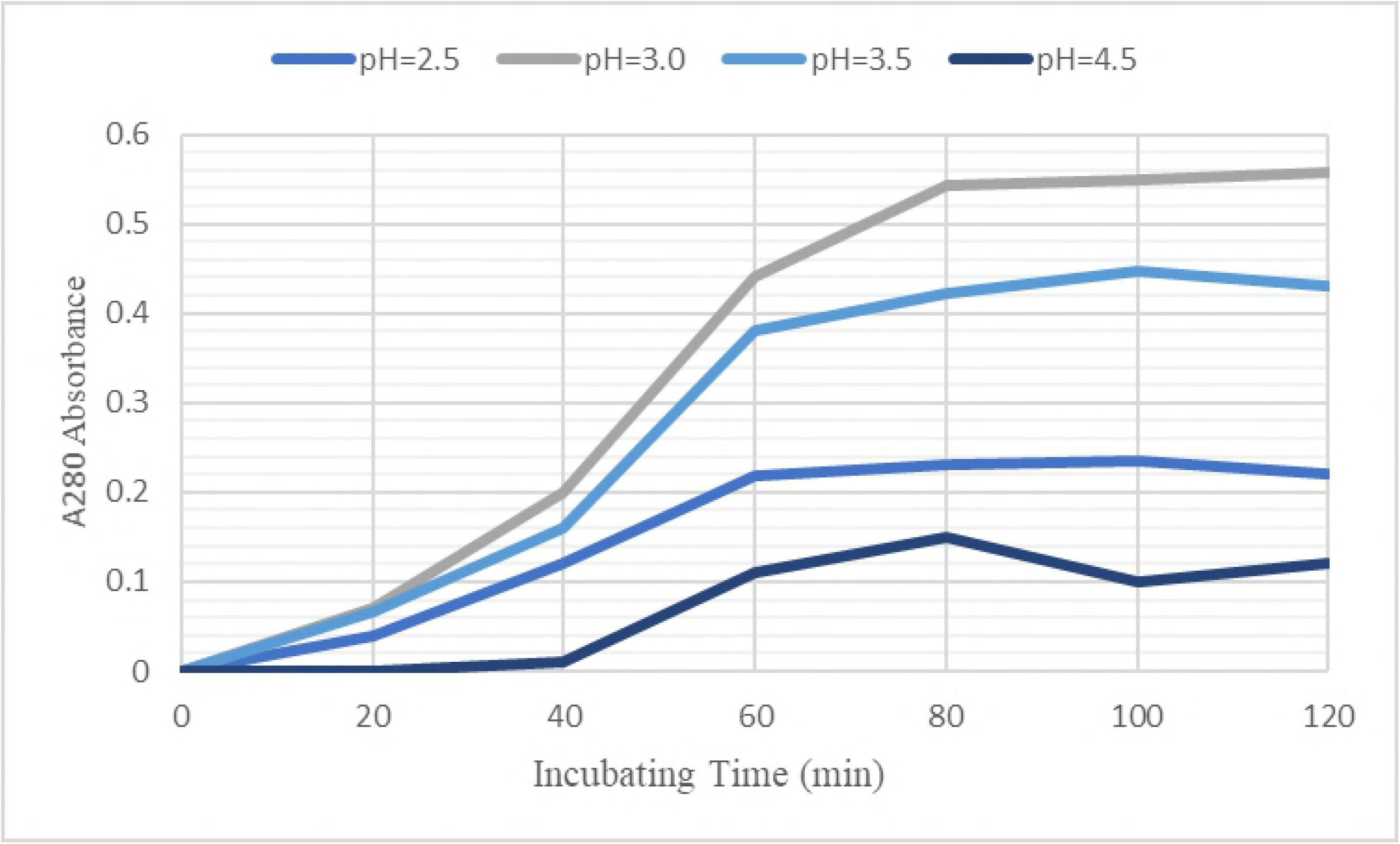
The results of the activated sjCTSD on the enzyme - cutting activity of bovine hemoglobin.

**Figure 8.**
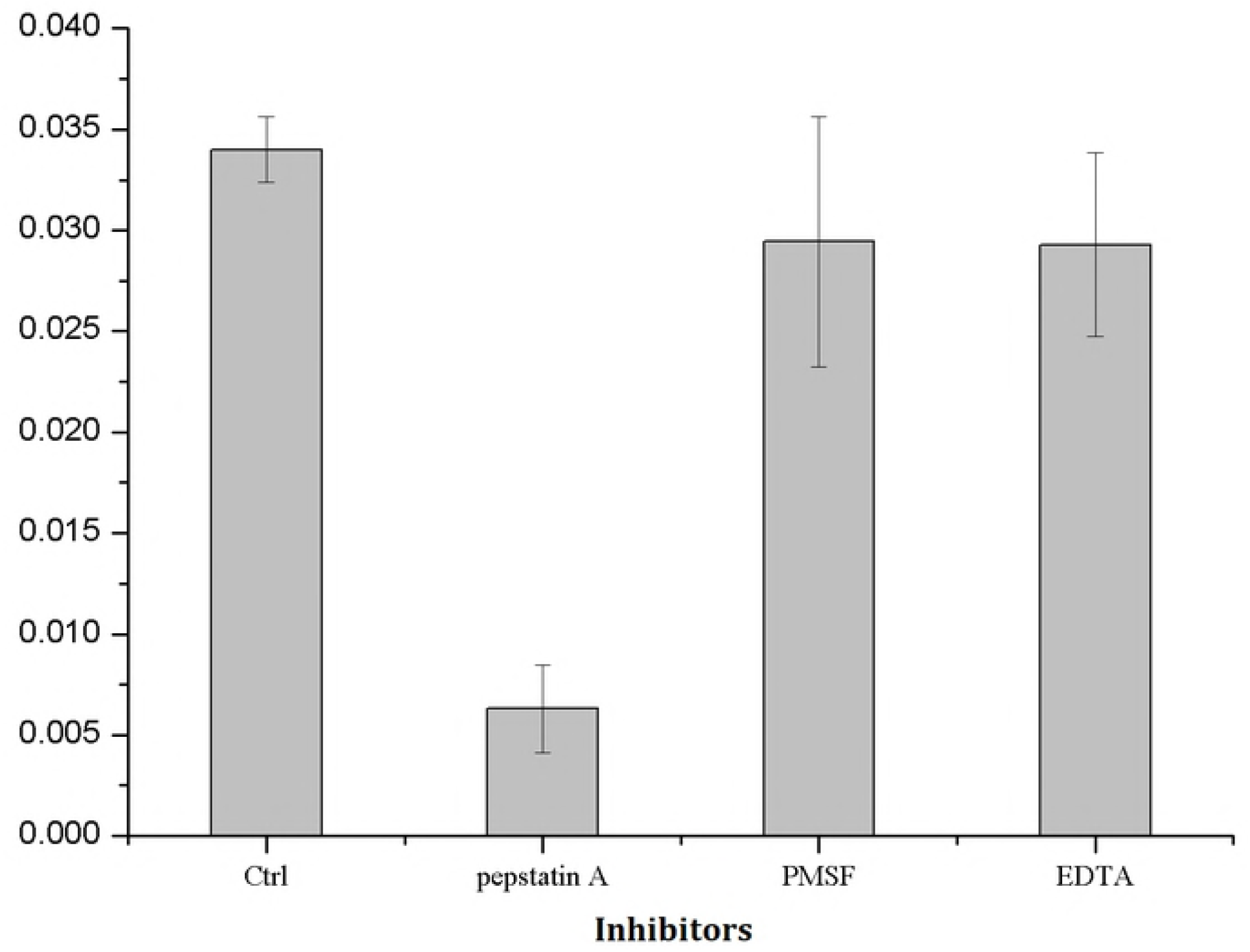
Effect of inhibitors on sjCTSD enzyme activity. Ctrl is the control group.

### 3.8 Antibacterial activity of sjCTSD

The difference of bacterial colonies was used as a marker to distinguish whether sjCTSD had a significant inhibitory effect on *E. coli* and *V. alginolyticus*. We calculated the survival rate of the colonies by comparing the experimental group treated with sjCTSD with the control group (survival rate = (control - experimental group)/control) (*p* < 0.05). As Table 2 shown that the activated sjCTSD has a significant inhibitory effect on the growth of *E. coli*, but no significant effect on the growth of *V. alginolyticus*.

**Table 2.** The antibacterial property of sjCTSD

| Protein concentration<br>($\mu\text{g/mL}$) | Clump count<br>(*E. coli*) | Survival rate<br>(*E. coli*) | Clump count<br>(*V. alginolyticus*) | Survival rate<br>(*V. alginolyticus*) |
| --- | --- | --- | --- | --- |
| 0 | 326 | 100% | 183 | 100% |
| 100 | 244 | 74.8% a | 169 | 92.3% |
| 200 | 190 | 58.3% a | 160 | 87.4% |
| 300 | 146 | 44.8% a | 151 | 82.5% |
| 400 | 153 | 46.9% a | 151 | 82.5% |

## 4. Discussion

The deduced amino acid sequence of CTSD of *S. japonica* was highly conserved with sequences of other invertebrates, especially *T. pacificus*. This study revealed that the sjCTSD protein had a molecular mass of 42.81 kDa and a theoretical pI of 5.52, similar with that of invertebrates (Komai et al., 2004; Pan et al., 2012). A previous study showed that the *Macrobrachium rosenbergii* CTSD gene encoded 385 amino acids (Zhang et al., 2016), while the sjCTSD encoded 393 amino acids. Previous studies showed that CTSD would be glycosylated and located transported to lysosomes through the endoplasmic reticulum (ER)-Golgi transfer pathway, after being synthesized in cytoplasm, and N-glycosylation plays an important role in this process. Through the multiple sequence alignment analysis with 12 other species, the first N-glycosylation site was more conserved among those species, the second N-glycosylation site was replaced by Asp in shrimps and *Polyrhachis vicina* (Cheng et al., 2014; Liang et al., 2010). The difference in character conservation may be related to the different functions of glycosylation.

Cathepsin D has a variety of biological functions, including intracellular and extracellular protein degradation, activation of zymogen, as a hormone, as a growth factor (Ohri et al., 2007), and is involved in apoptosis (Vashishta et al., 2006) etc. In the present study, analysis of the spatio-temporal expression profiles of the sjCTSD gene showed that the mRNA was expressed in all tissues in *S. japonica*. In addition, previous studies showed that CTSD is related to immune response, the expression of the shrimp CTSD gene in the hepatopancreas and hemolymph has been shown to significantly increase with the time of infection (Duan et al., 2013, Zhang et al., 2016b). The present study showed the expression levels of the sjCTSD gene continued to increase with the growth of *S. japonica*, suggesting that sjCTSD may participate in the aging process.

The expression level of sjCTSD activity was low during the early growth stages of the cuttlefish, but significantly increased during the late growth stages of *S. japonica*. From the young to prespawning stage, the level of sjCTSD was elevated because this stage represents a period of growth and maturation; during this time *S. japonica* undergoes fast physical development and the up-regulation of the sjCTSD gene may have resulted from individual morphological development and vigorous cell-replacement processes. This result was consistent with Saftig (1995), who found that mice stop growing after CTSD gene was knocked, thereby demonstrating the importance of the CTSD gene in the development of the species.

Previous studies indicated that the highest expression of cathepsin D in gonads was related to the role of this enzyme in yolk formation in *Sparus aurata* (Carnevali et al. 2001; 2003; 2008). It is consistent with our result that an increasing level of sjCTSD gene was detected in the prespawning stage, compared to the young stage. However, the expression in cuttlefish gonads require to be further examination.

Previous studies have shown that the pancreas and liver are the major digestive and metabolic organs, as well as the energy-hoarding organs of *S. japonica* (Fan et al. 2008). In addition, studies have shown that the brain, optic glands, and optic lobes of cuttlefish are important endocrine organs and the primary secretory tissue of gonadotropin-releasing hormone (GnRH; Yu 2011). Gonadotropin-releasing hormone and its analogues also has function in stimulating reproduction activity in invertebrates; injecting GnRH can induce spawning in some viviparous tunicates (Fiore et al. 2000; Adams et al. 2003; Chen et al. 2003). Once *S. japonica* enters the third growth stage, its physiology and metabolism start failing. In the postspawning stage before death, the sjCTSD expression levels were significantly increased in the optic glands, suggesting that the over-expression of sjCTSD had a negative effect on the regulatory function of optic glands. Degenerative optic glands can’t secrete GnRH to stimulate *S. japonica* to spawn and this result in many remnant eggs, which hinders the stable development of *S. japonica* eggs. Similarly, the sjCTSD expression level in the pancreas and liver both increased significantly. *S. japonica* may even stop feeding to protect eggs during the spawning stage, which is consistent with octopus (Wodinsky 1977). Expression of sjCTSD significantly increased from spawning onward, suggesting that sjCTSD promotes the release of energy stored in the liver and pancreas for spawning. Rapid depletion of energy store and small meals are both reasons for the mortality rate to skyrocket, so it was considered that sjCTSD also may play a potential role as a cuttlefish killer. Further research involved in gene knockout or editing is needed to understand the specific mechanisms involved.

Under the condition of 25°C incubation, the acidic buffer solution can induce the prokaryotic recombinant protein sjCTSD to self-cleave the N-terminal propeptide to reveal active sites. SjCTSD has the highest autolytic activity, when the pH value is 3.0. However, sjCTSD loses its ability to self-cut and cannot activate enzyme activity at neutral or alkaline buffer. Cathepsin D of *T. pacificus*, similar with sjCTSD, could gat the highest autolytic activity at pH=3.0 with the specific reaction substrate (Komai et al. 2004). *Eriocheir sinensis* is an important freshwater economic product in China, the enzyme activity of its CTSD can exert enzyme activity from pH 4 to 7 and achieve the highest at pH=5.0; The enzyme activity of *Euphausia superba* CTSD got the highest at pH=8.0, but the enzyme activity was significantly decreased in pH=9.0, the probably explanation was that the loss of activity in the alkaline environment (Zhu. 2017). In this paper, we observe that the optimal pHs for different species of CTSD were different. The species with closer homology, the physical and chemical properties of their CTSD are closer, however, different substrates specific for CTSD may also be responsible for the difference in the optimal pH because of the inconsistency of the experiments (Stoknes et al. 1995; Wang et al. 2007). CTSD is an aspartic protease that cannot be inhibited by serine protease inhibitors and metalloproteinase inhibitors. This Study have shown that recombinant CTSD expressed in prokaryotic vectors exist in the form of inclusion bodies. After destroying the existing structures of inclusion bodies, the denaturants that have a negative effect on protein conformation were removed and contribute to restoring the correct conformation of CTSD and subsequently recovering the enzymatic activity. The inhibition of the sjCTSD enzyme activity by the aspartic protease inhibitor pepstatin A indicates that sjCTSD had already possessed the basic properties of aspartic protease after renaturation.

## 5. Conclusion

In this study, a cDNA clone of CTSD (1392 bp long, encoding a 393 amino acid protein) was cloned and characterized from *S. japonica*. The CTSD transcript was widely distributed in body tissues. With the individual development, CTSD gene expression gradually increased; CTSD expression of the postspawning stage was significantly higher than young *S. japonica*. The results of the CTSD spatio - temporal expression profiling of optic glands suggested that sjCTSD played a crucial role in spawning through regulation of GnRH levels and the high expression level in the pancreas and liver suggested that sjCTSD had a potential function in the death mechanism of cuttlefish. The prokaryotic expression vector pET28a/sjCTSD was constructed and transfected into *E. coli*, and then induced by IPTG to achieve purified recombinant protein. Through the identification of specific substrates and protease inhibitors, sjCTSD has successed in expression and extraction. The renaturation-concentrated protein concentration was determined to be 0.69 mg/mL by the BCA protein concentration detection kit. The detection of the specific substrate digestive activity showed that sjCTSD had a certain enzymatic activity on bovine hemoglobin after renaturation. We also found that sjCTSD had a certain antibacterial activity in *E. coli*. However, further studies are required on the molecular mechanism of CTSD regulation of aging processes in *S. japonica*.

## Acknowledgments

This research was supported by the National Natural Science Foundation of China (31101937, 41406138, 41676153), Hong Kong, the Macao and Taiwan science and technology cooperation project (2014DFT30120), Zhejiang Provincial Natural Science Foundation of China (LY17C190006), National Spark Program (2015GA700014), Zhejiang science and technology Program (2015F50055).

